# Multiple acyl-CoA thioesterases occupy distinct functional niches within the mitochondrial matrix

**DOI:** 10.1101/705327

**Authors:** Carmen Bekeova, Lauren Anderson-Pullinger, Kevin Boye, Felix Boos, Yana Sharpadskaya, Johannes M. Herrmann, Erin L. Seifert

## Abstract

Acyl-CoA thioesterases (Acots) hydrolyze fatty acyl-CoA esters. Acots in the mitochondrial matrix are poised to mitigate β-oxidation overload that may contribute to lipotoxicity. Several Acots associate with mitochondria, but whether they all localize to the matrix, and are redundant or have different roles is unresolved. We compared mitochondrial Acots (Acot2, 7, 9, and 13) in terms of suborganellar localization, activity, expression and regulation, in mitochondria from multiple mouse tissues and from a new model of Acot2 depletion. Acot7, 9 and 13 localized to the matrix, joining Acot2 that was previously shown to localize there. Mitochondria from heart, skeletal muscle, brown adipose tissue and kidney robustly expressed Acot2, 9 and 13, though Acot9 was substantially higher in brown adipose tissue and kidney mitochondria, as was activity for C4:0-CoA, a unique substrate of Acot9. In all these tissues, Acot2 accounted for ∼half of the thioesterase activity for C14-CoA and C16:0-CoA. In contrast, liver mitochondria from fed and fasted mice expressed little Acot activity, and this activity was confined to long-chain CoAs, and due mainly to Acot7 and Acot13 activity. Matrix Acots occupied different functional niches, based on substrate specificity (Acot9 *vs*. Acot2 and 13) and strong CoA inhibition (Acot7, 9, 13 but not Acot2). Interpreting these results in the context of β-oxidation, CoA inhibition would prevent Acot-mediated suppression of β-oxidation while providing for an Acot-mediated release valve when CoA is limiting. This release valve would operate across a wide range of acyl-CoA chain lengths. In contrast, CoA-insensitive Acot2 could provide a constitutive syphon for long-chain fatty acyl-CoAs. These results reveal how the family of matrix Acots can help to mitigate β-oxidation overload and prevent a CoA limitation.

## INTRODUCTION

Acyl-CoA thioesterases (Acots) hydrolyze acyl-CoA into CoA and an acyl chain, and have been classified into two families based on functional domain. Type I Acots are members of the superfamily of α/β-hydrolases that includes lipases, appear to be devoid of other functional domains, and are found only in mammals. Humans have 4 type I Acots, and mice and rats have 6. Type I Acots have a high degree of homology and are thought to have arisen from gene duplication. Humans and rodents possess Acots localized to the cytoplasmic (Acot1), mitochondria (Acot2) and to peroxisomes (Acot3-6 in rodents, Acot3-4 in humans). In contrast, Type II Acots (Acot7-15) have homologs across multiple phylogeny and share little homology beyond a hotdog fold domain that is single or tandem depending on the Acot. All Type II Acots possess StAR-related lipid transfer domains, at least one of which was shown to regulate activity (1), and some Type II Acots also contain other functional domain(s) (2). Type II Acots are found in the cytoplasm (Acot7-14), mitochondria (Acot7-13, 15), and peroxisomes (Acot8). Dual localization is also a possibility (Acot7, 11, 13). Each of the Type I and II Acots has a signature substrate specificity that includes saturated and unsaturated fatty acyl-CoAs of different chain lengths and in fewer cases other CoA esters such as methyl branched-chain CoAs (for review: (1,3,4)).

Acots are predicted to have a high biological relevance because their substrates, acyl-CoA esters, are also substrates for other enzymes that serve major metabolic pathways, such as mitochondrial β-oxidation. Interestingly, β-oxidation overload has been described in the context of high substrate supply, and the bottleneck was localized to the mitochondrial matrix (5) raising the question of a role for Acots to mitigate the overload. Beyond β-oxidation, acyl-CoA esters can serve as allosteric or covalent regulators, which greatly expands the potential biological relevance of thioesterases. In fact, genetic manipulation in mice of Type I (Acot1 and Acot2) or Type II (Acot7, 11, 13 and 15) Acots has in all cases been associated with altered phenotypes (for review: (2,6,7)).

The recent availability of genetic mouse models for the Acots is sure to continue to provide information into the biological relevance of the Acots. Yet, there are basic unexplored questions about the Acots, the answers to which will help provide mechanistic insights into how Acots can alter cell phenotypes and also how to interpret models of depletion of one of the Acots. Fundamental considerations concern the localization (tissue, subcellular and suborganellar expression), regulation and the extent of thioesterase activity for particular substrates vis-à-vis the activity of major competing enzymes. What makes these considerations particularly important is the existence of so many Acots, some of which overlap in tissue expression, subcellular localization and substrate specificity. Thus a fundamental question is whether the Acots within a cellular compartment occupy functional or tissue-specific niches or whether, instead, the different Acots are largely redundant, with distinction only in substrate-specificity.

To investigate this question, we focused on the Acots that have been associated with mitochondria: Acot2, 7, 9, 11, 13 and 15. Indeed, it is already known that multiple Acots localize, or have the potential to localize, to mitochondria, and that mitochondria from some tissues express more than one Acot. Thus, some tissues have the potential to possess an abundance of mitochondrial thioesterase activity, and for this activity to originate from multiple Acots, which, superficially, would suggest redundancy among mitochondrial Acots; but whether this is the case has not been considered. In particular, what is not clear is the total Acot activity within mitochondria of different tissues, the substrate-specificity of this thioesterase activity within mitochondria from a given tissue, and the mitochondrial Acots that would contribute to this activity within a given tissue. These considerations would address the question of functional niche within a tissue as well as tissue-specific niches. Also, the sub-compartmentalization within mitochondria is not known for all of the Acots, and would further define niche *versus* redundancy. Finally, regulation is incompletely understood, and can also define functional niche.

We addressed these questions by first evaluating the suborganellar localization of Acot7, 9 and 13 which had not previously been determined. Then, to evaluate tissue- and substrate-specificity, we undertook a comparison of four mitochondrial thioesterases (Acot2, 7, 9, and 13) in terms of activity and expression, using five different mouse tissues. To compare among these Acots in terms of regulation by metabolic stimuli, effects on their expression and activity of fasting and potential allosteric regulators were evaluated in mitochondria from three mouse tissues and recombinant enzyme. Finally, using information from the above studies along with a new mouse model of Acot2 depletion, we estimated the quantitative contribution of Acot2 and Acot13 towards the total thioesterase activity for long-chain fatty acyl-CoA esters. All together these data reveal distinct functional roles for four mitochondrial Acots. Information from this study is also expected to facilitate the development of hypotheses for the biological roles of the Acots in different tissues, and how to study these hypotheses.

## RESULTS

### Localization of Acot7, 9 and 13 to the mitochondrial matrix

Sub-compartmental localization of Acot2 and Acot15 to the mitochondrial matrix was previously demonstrated by immuno-gold electron microscopy (8,9) whereas localization of Acot7, 9 and 13 within mitochondria has not been experimentally determined. Acot7 and 9 each express an N terminal mitochondrial targeting sequence (MTS) with high probability (84% or 93% for Acot7, depending on isoform, see below; 93% for Acot9; MTS predictions from TargetP). Acot13, with only a weak TargetP score for presence of an MTS (30%), has been suggested to reside on the outer surface of the outer mitochondrial membrane (OMM) (2), but this has not been demonstrated. Finally, whether Acot11, with only a low probability MTS prediction, localizes to mitochondria has not been clear and was also evaluated here. We used a protease protection approach to analyze the sub-compartmentalization of Acot7, 9 and 13 using mitochondria from different tissues. The ability of the protease to destabilize the protein upon strong membrane disruption by Triton X-100 served as a positive control for sensitivity to the protease. Tom20 and Tim23 were used to mark the OMM and IMS/IMM, respectively. PDH (pyruvate dehydrogenase kinase) marked the matrix. We used well-established antibodies for Tom20, Tim23 and PDH.

*Acot13*. The Acot13 antibody was shown to recognize a non-specific band at the predicted molecular weight for Acot13 (15 kDa) when tissue lysates from Acot13 knockout mice were analyzed (David Cohen, personal communication). Thus we determined the specificity of the antibody using recombinant Acot13 (**Fig1A**, left panel) and liver mitochondria from Acot13 knockout and control mice (**Fig1A**, right panel). Control samples revealed a single band at ∼15 kDa, whereas bands in this range were not detected in the Acot13 knockout sample unless the membrane was overexposed in which case faint bands were similar across protease/detergent conditions (**Fig1A**, right panel). Thus, the anti-Acot13 antibody is specific for Acot13 in isolated mitochondria.

**Figure 1.**
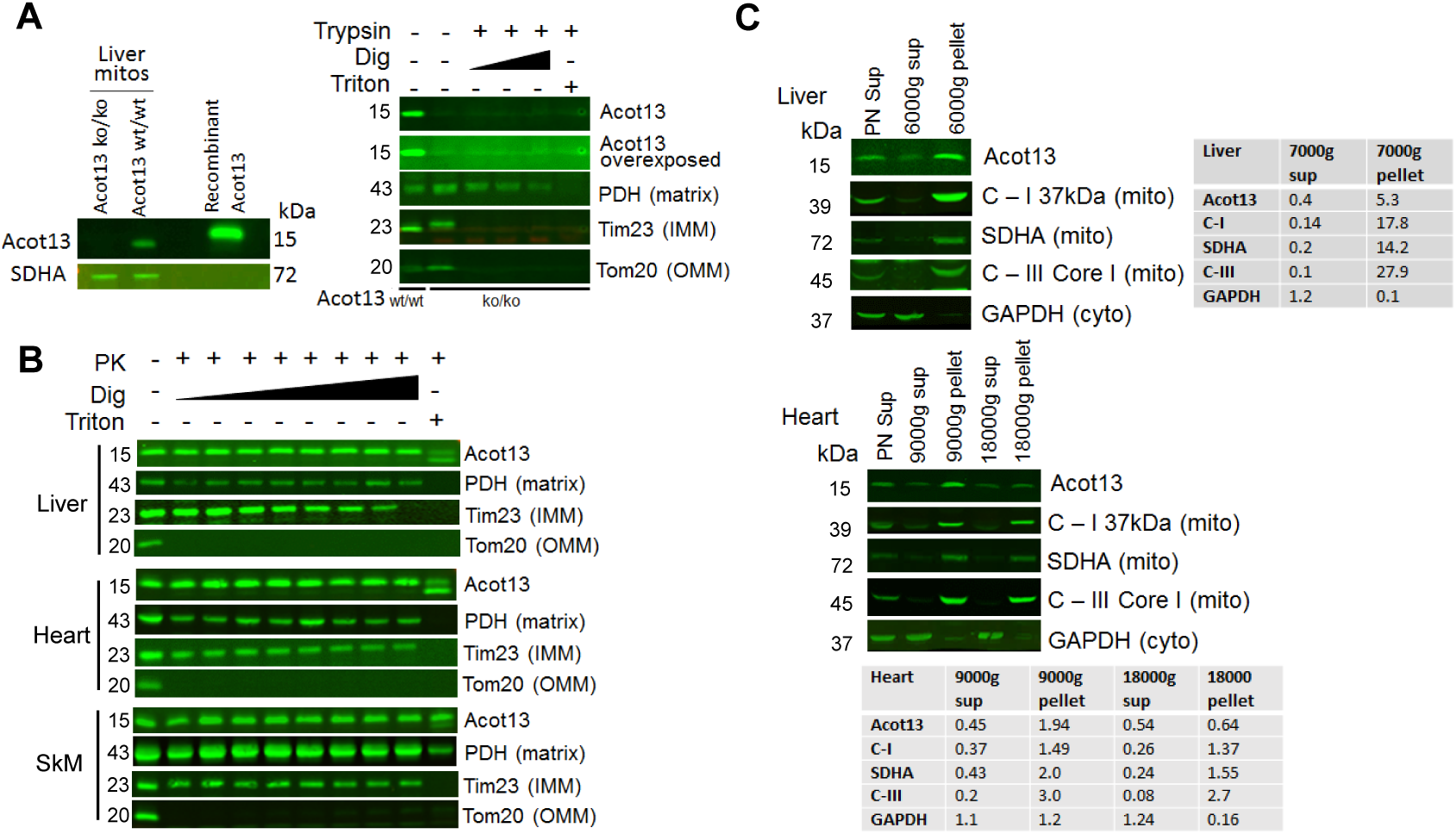
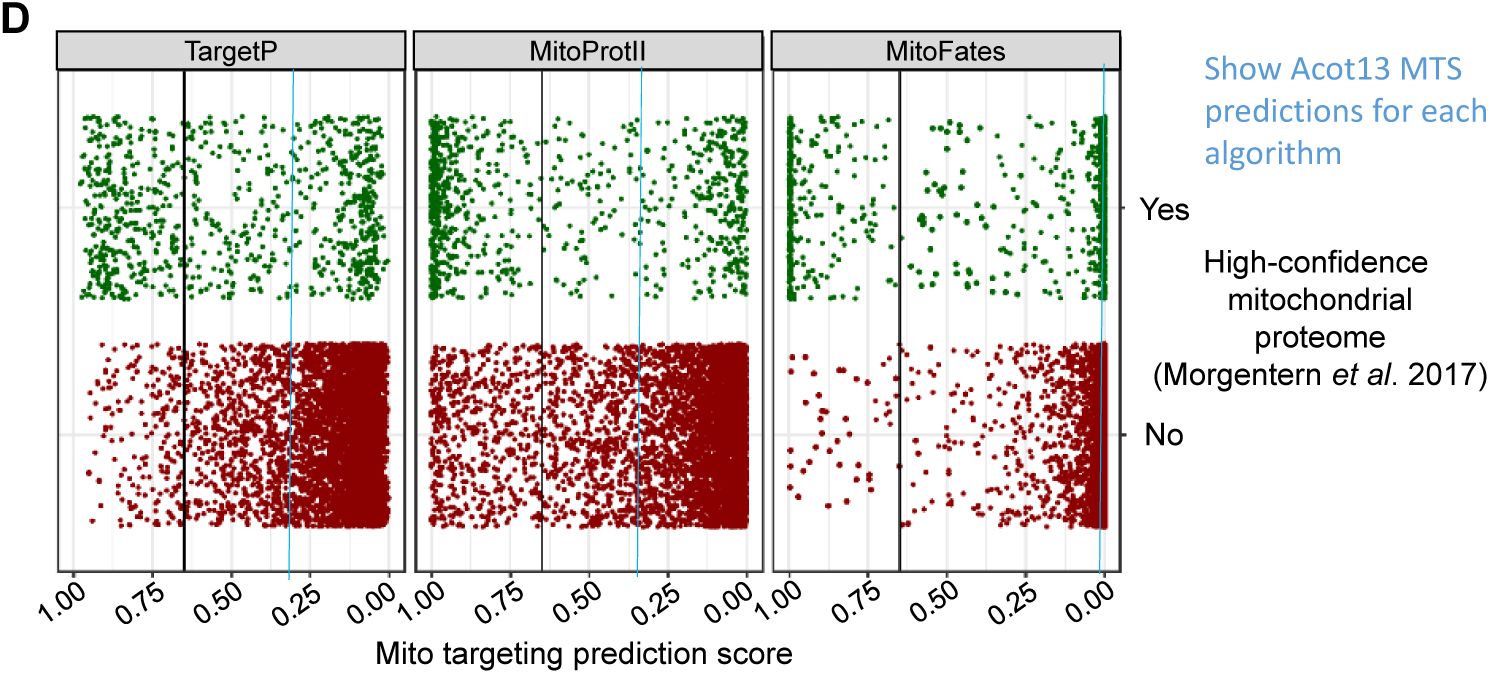
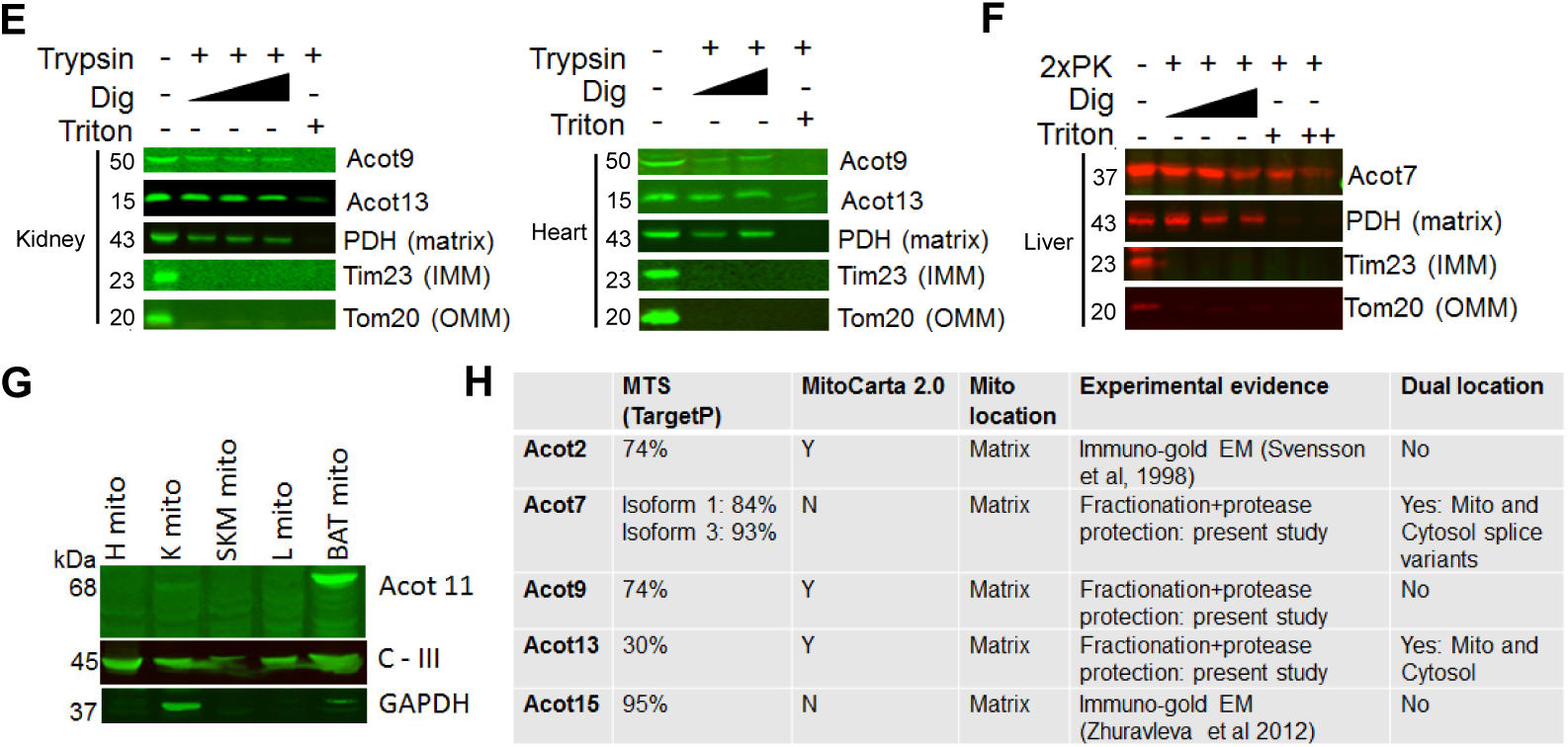
Acot7, 9 and 13 localize to the mitochondrial matrix. Protease protection experiments (**A, B, E, F**) determined the sub-mitochondrial location of Acot7, 9 and 13 using trypsin or proteinase K (PK), and increasing digitonin (Dig) concentration. Triton X-100 plus protease served as a positive control for protease sensitivity. Tom20 marked the outer mitochondrial membrane (OMM), Tim23 marked the intermembrane space/inner mitochondrial membrane (IMM) and pyruvate dehydrogenase (PDH) marked the matrix. **All panels**: mitochondria were isolated from the indicated mouse tissue. **A**: Recombinant Acot13 and mitochondria from Acot13 knockout (ko/ko) and wild type (wt) mice were used to determine the specificity of the anti-Acot13 antibody. **B**. Protease protection experiment to determine the sub-compartalization of Acot13 from the mitochondrial fraction of a liver fractionation. **C**: Distribution of Acot13 between mitochondria (pellet) and non-mitochondrial fraction. Subunits of electron transport chain Complex I, II and III (C-I, SDHA (succinate dehydrogenase)) were used to mark mitochondria. GADPH (glyceraldehyde 3-phosphate dehydrogenase) marked the cytoplasm. **D**. Mitochondrial targeting sequence (MTS) prediction accuracy by 3 algorithms using data from a high-confidence mitochondrial proteome from yeast (11). Yeast proteins were classified as mitochondrial or non-mitochondrial according to (11). The amino acid sequences of the complete proteome of reference strain S288C were retrieved from the Saccharomyces Genome Database and subjected to MTS prediction using the TargetP, MitoProt II and MitoFates algorithms. The resulting prediction scores for mitochondrial localization are plotted as points for each individual protein. **E**. Protease protection experiment to determine the subcompartalization of Acot13 present in the mitochondrial fraction of kidney and heart fractionations. **F**: Acot11 is not widely expressed in mitochondria. H: heart, K: kidney, SKM: skeletal muscle, L: liver, BAT: brown adipose tissue. **G**: Summary of mitochondrial Acots, likelihood of a mitochondrial targeting sequence (MTS, TargetP algorithm), and localization within mitochondria.

We next exposed mitochondria isolated from three different tissues to each of two different proteases in the absence or presence of Triton X-100 or increasing digitonin, then evaluated the expression of Tom20, Tim23, PDH and Acot13. Tom20 was digested in the presence of protease alone, Tim23 usually required addition of digitonin for digestion and was more susceptible to trypsin than to proteinase K, and PDH was only digested when Triton X-100 was added (**Fig1A, B, E, F**). In mitochondria from all three tissues (liver, kidney and heart) and using either trypsin or proteinase K, Acot13 was only digested in the presence of Triton X-100 and thus followed the pattern of matrix-localized PDH (**Fig1B, E**).

It was formally possible that some Acot13 was loosely associated with the outer surface of the OMM but then was dislodged during tissue homogenization. We reasoned that if this occurred, the supernatant fraction of the high speed spin used to pellet mitochondria would contain a substantial amount of Acot13. Indeed, Acot13 was previously found in the supernatant fraction and it was concluded that it can reside in the cytoplasm as well as be associated with mitochondria (1). To estimate the fraction of Acot13 in the cytoplasm (or loosely bound to the OMM), immunoblot analysis was conducted on different fractions from the mitochondrial isolation, namely the total homogenate prior to centrifugation, the supernatant from the first high speed spin, and the washed pellet from the final high speed spin. Liver and heart mitochondria were tested (**Fig1C**). Compared to the amount of Acot13 detected in the total homogenate, 5x more was found in the high speed pellet from liver and ∼2x more was found in the high speed pellet from heart. In contrast, band intensity in the supernatant of both tissues was less than half of that in the total homogenate. For the heart, this pattern of distribution of Acot13 between pellet and supernatant was similar to that for three known inner mitochondrial membrane (IMM) proteins (Complex I 37 kDa subunit, Complex II SDHA subunit, Complex III Core I subunit), raising the possibility that Acot13 detected in the supernatant reflects contaminating mitochondrial proteins rather than a cytoplasmic or OMM-associated fraction.

The situation for the liver was different. In the liver, Acot13 in the supernatant as a fraction of the total homogenate was substantially higher than that for any of the three IMM proteins (0.4 for Acot13 *vs*. 0.1-0.2 for C-I, SDHA and C-III), suggesting that, in the liver, some Acot13 was present in the cytoplasm and/or very loosely associated with the OMM. Also, compared to the three IMM proteins, there was less enrichment of Acot13 in the high speed pellet whereas in the heart, enrichment of the Acot13 and the IMM proteins was more similar. However, for both liver and heart, the majority of Acot13 was present in the high speed pellet and thus would situate a substantial amount of the Acot13 within the mitochondrial matrix.

Analysis using MTS prediction algorithms indicated a probability of an MTS in mouse Acot13 of ∼30% (TargetP algorithm) or 39% (MitoProt II algorithm; cleavage site could not be predicted). MTS prediction for human Acot13 isoform 1 was higher (∼52% using TargetP; 63% using MitoProt II; cleavage site not predictable), and was much lower for human Acot13 isoform 2 (∼12% using TargetP; 50% using MitoProt II and with a predicted cleavage site at amino acid 13). We searched for an internal targeting sequence (10) by sequentially shortening the amino acid sequence by 1 amino acid starting at the N terminal. This analysis failed to reveal an MTS with a probability higher than 20% anywhere within the protein.

Perusing MitoCarta2.0 revealed *bona fide* mitochondrial matrix proteins having MTS predictions <30% (e.g., ribosomal proteins MRPS22, MRPL42 and MRPL48). This, together with our matrix localization of Acot13 despite its low MTS prediction score prompted us to systematically determine the reliability of MTS algorithms. For this we used a recently published high-confidence mitochondrial yeast proteome (11). Using the TargetP, MitoFates and MitoProt II prediction algorithms, and a cutoff MTS probability of 65%, the false negative rate (high-confidence mitochondrial proteins with MTS < 65%) was 60%, 56% and 38% for each algorithm respectively. The false positive rate (high-confidence non-mitochondrial protein with MTS>65%) was 4%, 1% and 14% for each algorithm respectively. Furthermore, plotting the MTS scores for high-confidence mitochondrial and non-mitochondrial proteins revealed many high-confidence mitochondrial proteins with MTS scores <25% (**Fig1D**). We also analyzed the human and mouse MitoCarta2.0 databases (12) for proteins annotated as “Apex_matrix”; ∼6% of these proteins had a low MTS score (category “5” in MitoCarta) or were categorized as having no MTS (“no score” in MitoCarta). These analyses provide a quantitative argument that MTS scores > 65% predict mitochondrial proteins well whereas a low MTS score is unreliable as evidence that a protein is non-mitochondrial. This can in part be explained by the fact that a number of mitochondrial proteins carry targeting signals other than a classical N-terminal MTS. While these are characterized for some classes of proteins (e.g., substrates of the TIM22 or MIA pathways), the targeting information for many of these proteins is unknown.

*Acot9*. Localization of Acot9 was tested in kidney mitochondria because they robustly express Acot9 (4)(see also **Fig2**). Heart mitochondria were also tested. A custom antibody tested in mouse tissues (4) was used that displays a band near 50 kDa, with minimal or no additional bands. In both kidney and heart mitochondria, the 50 kDa band disappeared only after adding trypsin plus Triton X-100, as was found for the matrix protein PDH (**Fig1E**).

**Figure 2.**
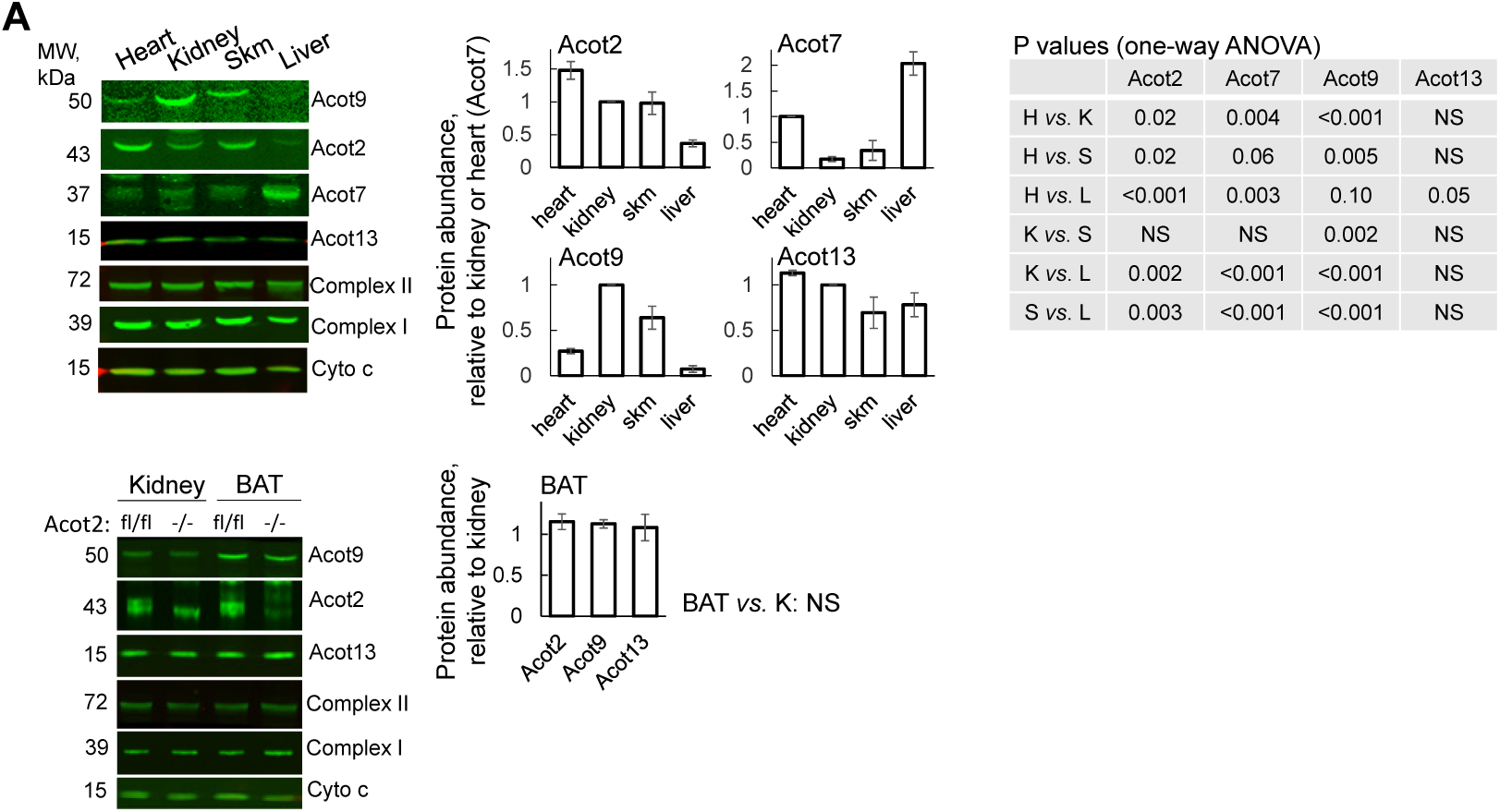
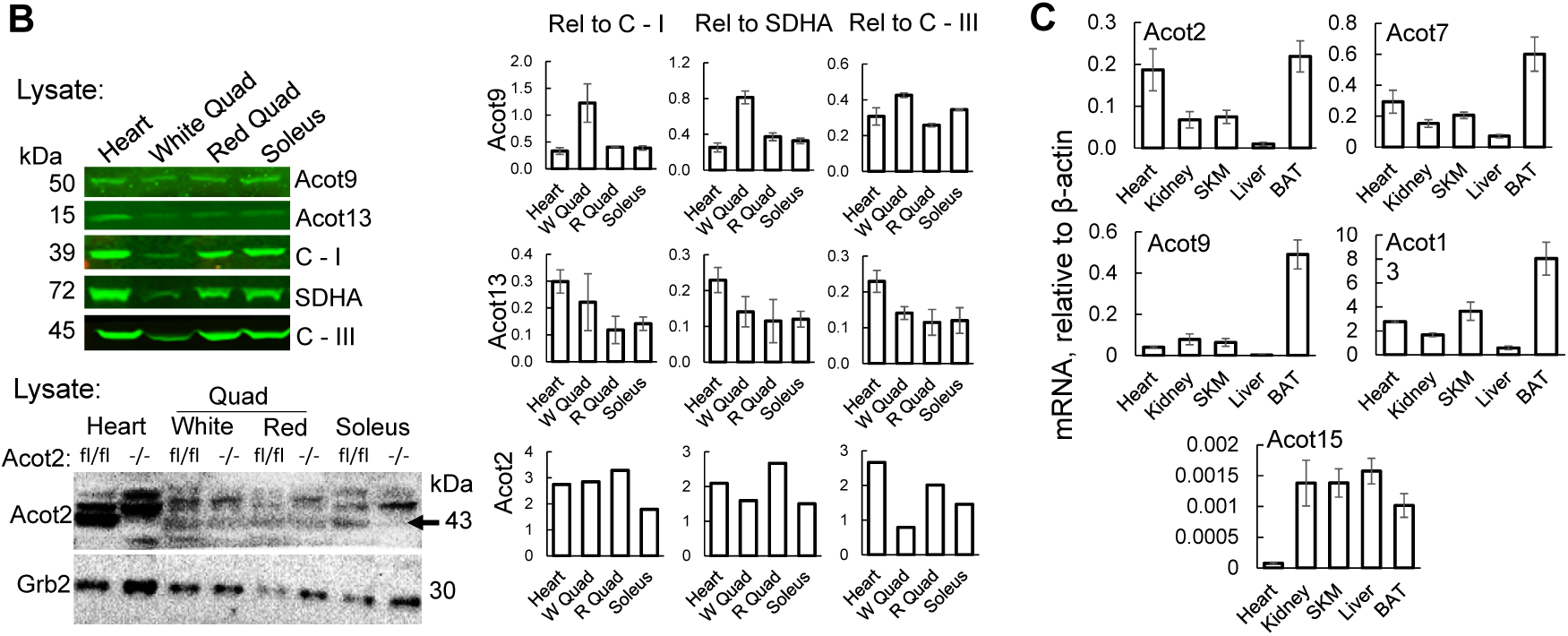
Tissue-specific localization of mitochondrial Acots. **A, B**: Immunoblot analysis using isolated mitochondria (**A**) or lysates from the heart and different muscles (Sol: soleus; W and R Quad: white and red quadriceps). Skm: skeletal muscle (**B**). In **A**, normalization was to Complex I, Complex II (SDHA: succinate dehydrogenase) and cytochrome C (cyto c); normalizations to each of the latter were averaged, then expressed relative to that for kidney or heart mitochondria (equal to 1.0). In **A**, the antibody against Acot2 used with the BAT samples yielded non-specific bands slightly below the Acot2 band. To interpret the Acot2 bands, mitochondria from Acot2^-/-^ tissues were run in parallel. Only the upper band of the wild-type samples was used for quantification. The Acot7 signal was not evaluated in BAT because the antibody yielded many non-specific bands. Statistics: One-way ANOVA, Tukey post hoc tests. **: p<0.05 Liver *vs*. all other organs; *: p<0.05 *vs*. Heart and Liver; #: p<0.05 Liver *vs*. Skeletal muscle, Heart and Kidney; $: p<0.05 Skeletal muscle and Liver *vs*. BAT and Kidney. **B**, the Acot2 antibody also recognizes Acot1 in the muscle lysates, thus the signal from tissue with loss of Acot2 (-/-) was taken as background and subtracted from the signal from tissue from floxed mice (fl/fl). **C**: mRNA expression. Values: mean±sem, n=5-6.

*Acot7*. Acot7 has three transcripts annotated in NCBI. Transcripts 1 and 3 each have a strongly predicted MTS (84% and 93%, respectively, using TargetP) whereas prediction for transcript 2 is 11%. For all transcripts, the molecular weight of the unprocessed protein is ∼42 kDa. Cleavage of an N terminal sequence in transcripts 1 and 3 sites yields a ∼37 kDa protein. The anti-Acot7 antibody was custom made and previously tested in tissues from Acot7 knockout mice (13). Using this antibody we found the predominant band to be at ∼37 kDa. Occasionally faint bands appeared at ∼42 kDa and ∼50 kDa. Our studies showed that liver mitochondria clearly express the 37 kDa band whereas signals were much weaker or barely detectable in mitochondria from other tissues (see **Fig2A**). Thus we selected only liver mitochondria for protease experiments. Similar to what we observed for Acot13 and Acot9, the 37 kDa band recognized by the anti-Acot7 antibody was destabilized only when the sample was treated with protease plus Triton X-100 (**Fig1F**), and thus followed the same pattern as matrix-localized PDH.

Finally, using an antibody against Acot11 validated in knockout tissue (14), a band was observed in the high speed pellet fraction in the BAT preparation but not in the high-speed pellet fraction from heart, skeletal muscle, liver or kidney (**Fig1G**).

In summary, the observations in mitochondria from multiple tissues point to a matrix localization for Acot7, 9 and 13. Furthermore, at least in liver, Acot13 resides to some extent in the cytoplasm or loosely associated with the outer OMM, but the majority is found in the matrix. Finally, Acot11 was not found in mitochondria from the tissues we tested except BAT. **Fig1H** summarizes the localization of the mitochondrial Acots determined here and in previous studies.

### Tissue-specific Acot expression and thioesterase activity

The above observations indicate that Acot13 joins Acot2, 7, 9 and 15 as an Acot that localizes to the mitochondrial matrix, raising the possibility of redundancy among 5 mitochondrial Acots. Tissue-specificity of expression would be one way that these Acots are not redundant. Some information about tissue expression is available. However this partial information is distributed among several studies making it difficult to compare the potential importance of each Acot within or among tissues. Here we undertook a comparison of Acot protein expression and thioesterase activity among mitochondria from 5 tissues. Two of these, liver and heart, have already been associated with mitochondrial Acot expression because of biological relevance or because of high inducibility of an Acot in that tissue (3,15-19). Three other tissues, skeletal muscle, brown adipose tissue (BAT) and kidney, are known to express mitochondrial Acots (4,13,18,20,21) but have been less studied. All tissues robustly oxidized fatty acids in mitochondria.

There were several methodological considerations for the immunoblotting. First, concerning the antibodies, those against Acot2, 7 and 13 have been tested in samples from knockout mice (9,13,22)(this study), whereas there is no reliable and available antibody against Acot15. Second, mitochondrial fractions were largely clear of cytoplasmic proteins (e.g., see **Fig1C**). Third, for normalization, samples were probed for 3 different proteins: the 37 kDa subunit of C-I, SDHA, and cytochrome c (cyto c); the average of the normalized values was used. A new aliquot of the Acot2 antibody that was used for BAT mitochondria generated multiple bands; for interpretation, these immunoblots included samples from Acot2 whole-body knockout mice (Acot2^-/-^, see **Fig3B**). BAT was excluded from the Acot7 analysis because the Acot7 antibody generated multiple bands around 37 kDa.

**Figure 3.**
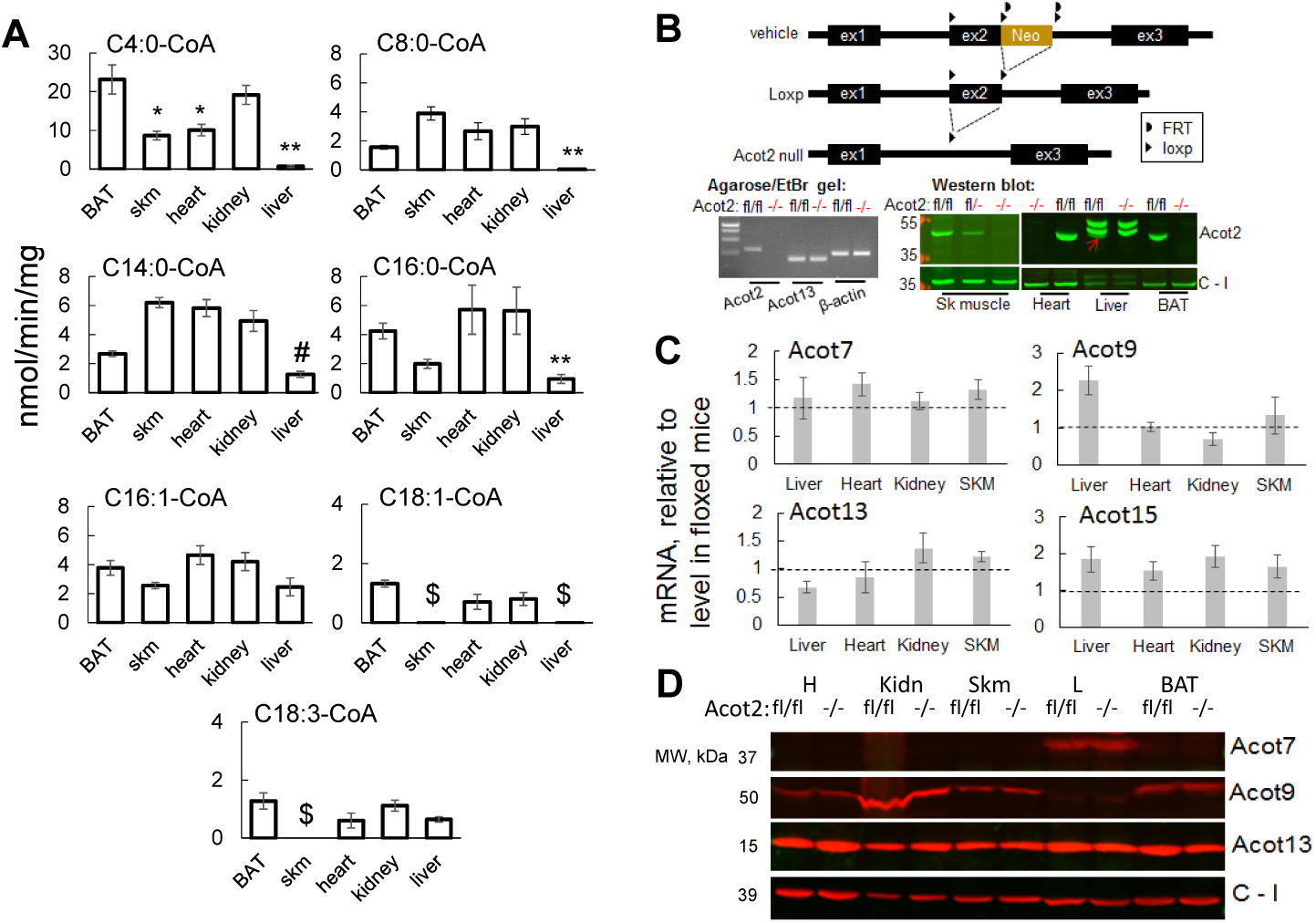
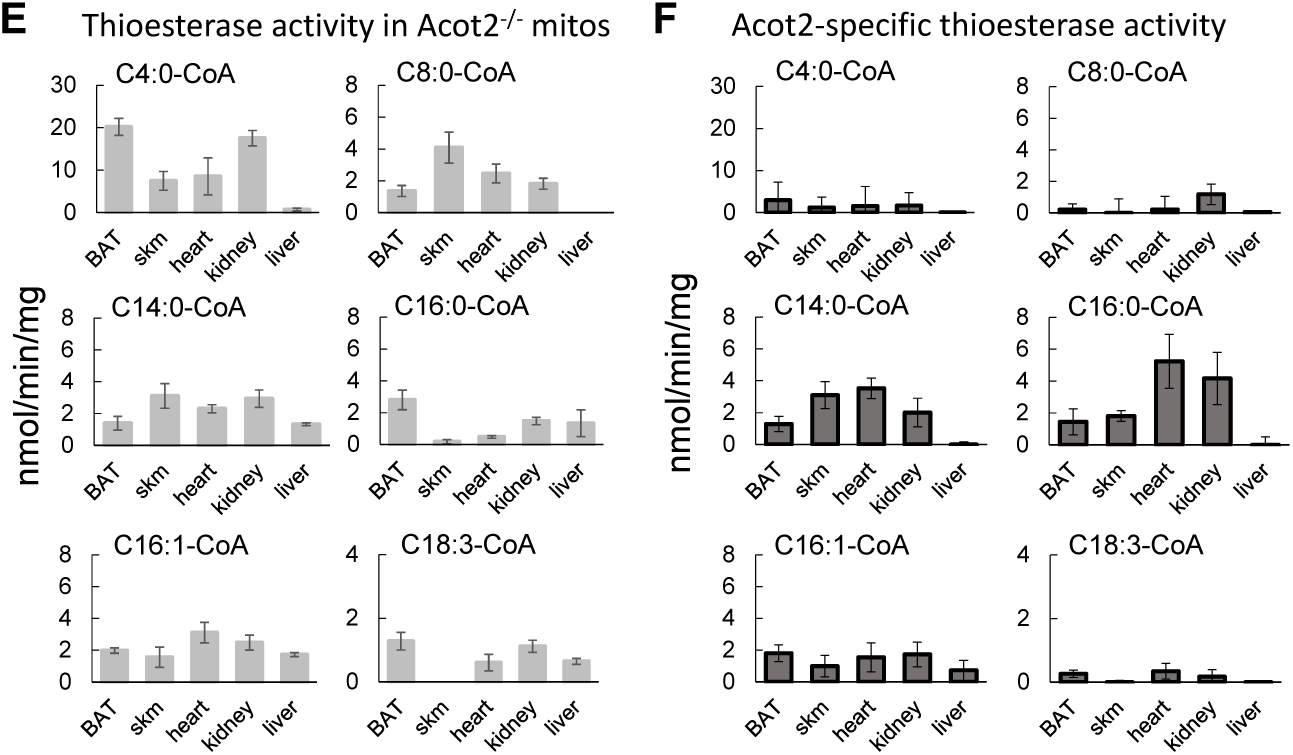
Tissue-specific mitochondrial thioesterase activity and contribution of Acot2. **A**: Thioesterase activity in mitochondria from mouse tissues (BAT: brown adipose tissue; skm: skeletal muscle) for a range of acyl-CoAs. Values: mean±sem; n=6-8. Table: p values, one-way ANOVA, Tukey post hoc tests; H: heart mitochondria; K: kidney mitochondria; S: skeletal muscle mitochondria; L: liver mitochondria; NS: not significant. **B**: Acot2 whole-body knockout mouse (Acot2^-/-^) model generated by crossing mice with floxed Acot2 alleles (fl/fl) with mice expressing Cre recombinase driven by the eIIa promoter. Knockout was validated by PCR and western blot (wb) analysis using isolated mitochondria. Arrow: faint Acot2 band in fl/fl liver mitochondria. **C, D**: Evaluation of possible compensatory changes in the expression of other mitochondrial Acots in Acot2^-/-^ mice at the level of mRNA (**C**) and protein (**D**). **E**: Thioesterase activity in mitochondria from Acot2^-/-^ mice. Values: mean±sem; n=4-5. **F**: Acot2-specific thioesterase activity calculated by subtracting activity measured in Acot2^-/-^ mitochondria (**E**) from activity measured in mitochondria from floxed mice (**A**). Error bars in (**F**) are the Gaussian error propagation (calculated as the square root of the sum of the variances).

Heart, skeletal muscle, BAT and kidney mitochondria all robustly expressed Acot2 and Acot13 (**Fig2A**). Skeletal muscle, BAT and kidney mitochondria also expressed Acot9, and heart slightly less so. Acot7 was low in mitochondria from these tissues. Liver mitochondria expressed Acot7 and Acot13, but very little Acot2 and Acot9. Thus mitochondria from skeletal muscle, BAT, kidney and heart robustly expressed 3 different matrix Acots (Acot2, Acot9, Acot13), whereas liver mitochondria show lower Acot expression, mainly contributed by Acot13 and Acot7.

Glycolytic and oxidative muscle fibers express a different abundance of proteins mediating metabolism and signaling, including proteins encoded by peroxisomal proliferator-activated receptor (PPAR)-induced genes. Because some Acots are induced by PPAR agonists (1,8,15,17,18,23-25), it was of interest to test Acot expression in different skeletal muscles. We evaluated the abundance of mitochondrial Acots in soleus (highly oxidative), white quadriceps (glycolytic) and the redder portion of gastrocnemius (mixed oxidative). Lysates were used because mitochondria cannot be reliably isolated from small pieces of mouse skeletal muscle. SDHA, C-I and cyto c levels were used to control for mitochondrial content differences. Because the Acot2 antibody also recognizes cytoplasmic Acot1, samples from Acot2^-/-^ mice were tested in parallel; the signal in Acot2^-/-^ muscle was taken as background and subtracted from wild-type samples. Acot9 showed relatively higher expression in white gastrocnemius compared to the oxidative muscles and the heart (**Fig2B)**. Differently, levels of Acot2 and Acot13 was similar in glycolytic and oxidative muscles (**Fig2B**).

Though a tissue comparison of Acot mRNA was recently published (13) as a complement to our dataset, we also evaluated mRNA levels in the 5 tissues studied here (**Fig2C**). Our data generally agree with published values. Notably, and similar to protein expression, Acot2, Acot7 and Acot13 mRNA were robust in all tissues except the liver.

As an orthogonal approach for understanding the relative contribution of different mitochondrial matrix Acots to a tissue’s thioesterase activity, we measured thioesterase activity using a range of acyl-CoA species that covers the substrate specificity of the matrix Acots. Activity measurements also allow quantitative comparisons among substrates within a tissue.

Thioesterase activity for C4:0-CoA was by far the highest, at 10-20 nmol/min/mg *vs*. 1-6 nmol/min/mg for the other substrates (**Fig3A**)(in all cases, unless otherwise stated, mg refers to mg of mitochondrial protein). BAT and kidney mitochondria showed the highest activity for C4:0-CoA (∼20 nmol/min/mg), followed by heart and skeletal muscle mitochondria (∼10 nmol/min/mg). Activity towards C4:0-CoA was very low in liver mitochondria. This pattern of activity for C4:0-CoA is fully consistent with the pattern of Acot9 protein abundance in these tissues and the fact that the only Acot known to show activity for C4:0-CoA is Acot9 (1,4,9,26).

Thioesterase activity for C8:0-CoA was lower (2-4 nmol/min/mg) than for C4:0-CoA but showed a similar tissue distribution including notable absence in liver mitochondria (**Fig3A**). As for C4:0-CoA, C8:0-CoA is a good substrate for Acot9 (4). Like C4:0-CoA, C8:0-CoA is not a good substrate for Acot2, 13 or 15 (1,8,9). Thus thioesterase activity towards C4:0-CoA and C8:0-CoA can be used as surrogate for Acot9 expression in BAT, kidney, heart and skeletal muscle.

Thioesterase activity for C14:0-, C16:0- and C16:1-CoA was 2–6 nmol/min/mg in mitochondria from all tissues, though activity in liver mitochondria was at the lower end of the range and activity in heart at the higher end (**Fig3A**). This pattern of activity among the tissues generally matched the protein levels of Acot2 and Acot13, both of which have C14:0-, C16:0- and C16:1-CoA as major substrates. Acot9 is reported to have substantial activity for C14:0-CoA (but not for Acot16:0- or Acot16:1-CoA) (4). Yet, activity for C14:0-CoA was not higher in BAT and kidney mitochondria which express the most Acot9. Similarly, C14:0-CoA and C16:0-CoA are good substrates for Acot7 (26), yet liver mitochondria, which expressed the most Acot7 among the 5 tissues, showed relatively low activity for these substrates. Thus Acot7 and 9 may show different substrate selectivity in their native context.

Finally we tested thioesterase activity for C18:1-CoA and C18:3-CoA, both of which are major substrates for Acot15 but poor substrates for the other mitochondrial Acots (4,9,24,26). In mitochondria from all tissues, activity for either substrate was 1-2 nmol/min/mg (**Fig3A**). Thus, the contribution of Acot15 to mitochondrial thioesterase activity is likely to be minor.

A question that arises is how much thioesterase activity within mitochondria is attributable to each Acot. An estimate for Acot9 activity in BAT, kidney and liver mitochondria was obtained using a C14:0-CoA thioether to inhibit Acot9 activity (4). Here we used a new mouse model of Acot2 loss to estimate the contribution of Acot2 to the total mitochondrial thioesterase activity. We generated mice with floxed Acot2 alleles (Acot2^fl/fl^), then used them to generate germline whole-body Acot2 knockout (Acot2^-/-^) mice (**Fig3B**). Mitochondria from these mice and floxed controls were used to estimate Acot2-specific activity. First, the possibility of upregulation of other mitochondrial Acots was considered. Acot2^-/-^ mice had higher mRNA for Acot9 in liver and Acot13 in kidney and skeletal muscle (**Fig3C**), but protein levels were not changed (**Fig3D**). Acot15 mRNA was higher in all Acot2^-/-^ tissues (**Fig3C**), but protein is unlikely to be higher because upregulated mRNA was still very low and thioesterase activity for its major substrate, C18:3-CoA, was not higher (see **Fig3E**).

**Fig3E** depicts thioesterase activity in all 5 tissues from Acot2^-/-^ mice. Acot2-specific activity was estimated as the difference in the average activity from mitochondria from control mice and from Acot2^-/-^ mice (**Fig3F**). Values for C4:0-CoA and C8:0-CoA in the Acot2^-/-^ samples were within the variability of the control samples (see **Fig3A**). Thus Acot2^-/-^ samples only showed activity for C14:0-CoA, C16:0-CoA and C16:1-CoA, and very low activity for C18:3-CoA, consistent with the expected substrate specificity for Acot2 (8). Acot2-specific thioesterase activity for these substrates in heart, skeletal muscle, kidney and BAT ranged from 2-5 nmol/min/mg. Liver mitochondria expressed very little Acot2-specific activity.

In the Acot2^-/-^ samples, activity towards C16:0-CoA and C16:1-CoA in heart, skeletal muscle, BAT and kidney likely reflects activity of only Acot13 because Acot7 is minimally expressed in those tissues and Acot9 has minimal activity for those substrates. Acot13 activity can be estimated as 2-3 nmol/min/mg towards C16:1-CoA and ∼1 nmol/min/mg towards C16:0-CoA except in BAT where it is ∼ 3 nmol/min/mg.

Overall, measurements shown in **Figure 3** indicate that the highest thioesterase activity in mitochondria is towards C4:0-CoA, catalyzed by Acot9, and is 3-10 times higher than activity towards C14:0-, C16:0- and C16:1-CoA which are catalyzed to a similar extent by Acot2 and Acot13. Activity towards C18:1- and C18:3-CoA is very low, suggesting only a minor contribution of Acot15 to mitochondrial thioesterase activity. Finally, BAT, kidney, heart, skeletal muscle mitochondria expressed far more thioesterase activity then liver mitochondria.

### Regulation of mitochondrial Acot protein abundance and activity

Some Acots were shown to be transcriptionally regulated by PPAR agonists and nutritional stimuli such as high fat feeding and fasting; studies have mainly shown this at the mRNA level (1,15,17,25,27), but some also at the protein level (8,17,23,28). Furthermore, Acot2 mRNA in the mouse heart oscillates with the time of the day, with the lowest expression near the end of the daily light phase (25). Thus, mitochondrial Acot activity may be more prevalent at certain times of the day due to time-dependent effects on gene regulation. To test this, we measured Acot protein abundance and thioesterase activity in mitochondria from liver, skeletal muscle and kidney from overnight fasted mice and compared it to that from mice ∼2 hrs before the end of the light phase. With fasting, Acot2 protein levels changed (∼20% rise) whereas protein levels of Acot9, Acot7 and Acot13 did not (**Fig4A**, quantified in **Fig4B**). Likewise, thioesterase activity in mitochondria from fasted mice was only increased for substrates that are major substrates for Acot2 (**Fig4C**). The overnight fasted condition likely represents an exacerbation of metabolic conditions during the mouse’s light phase. Thus, although some of the Acots are readily inducible by PPAR agonists and high fat feeding, inducibility by physiological stimuli appears to be milder.

**Figure 4.**
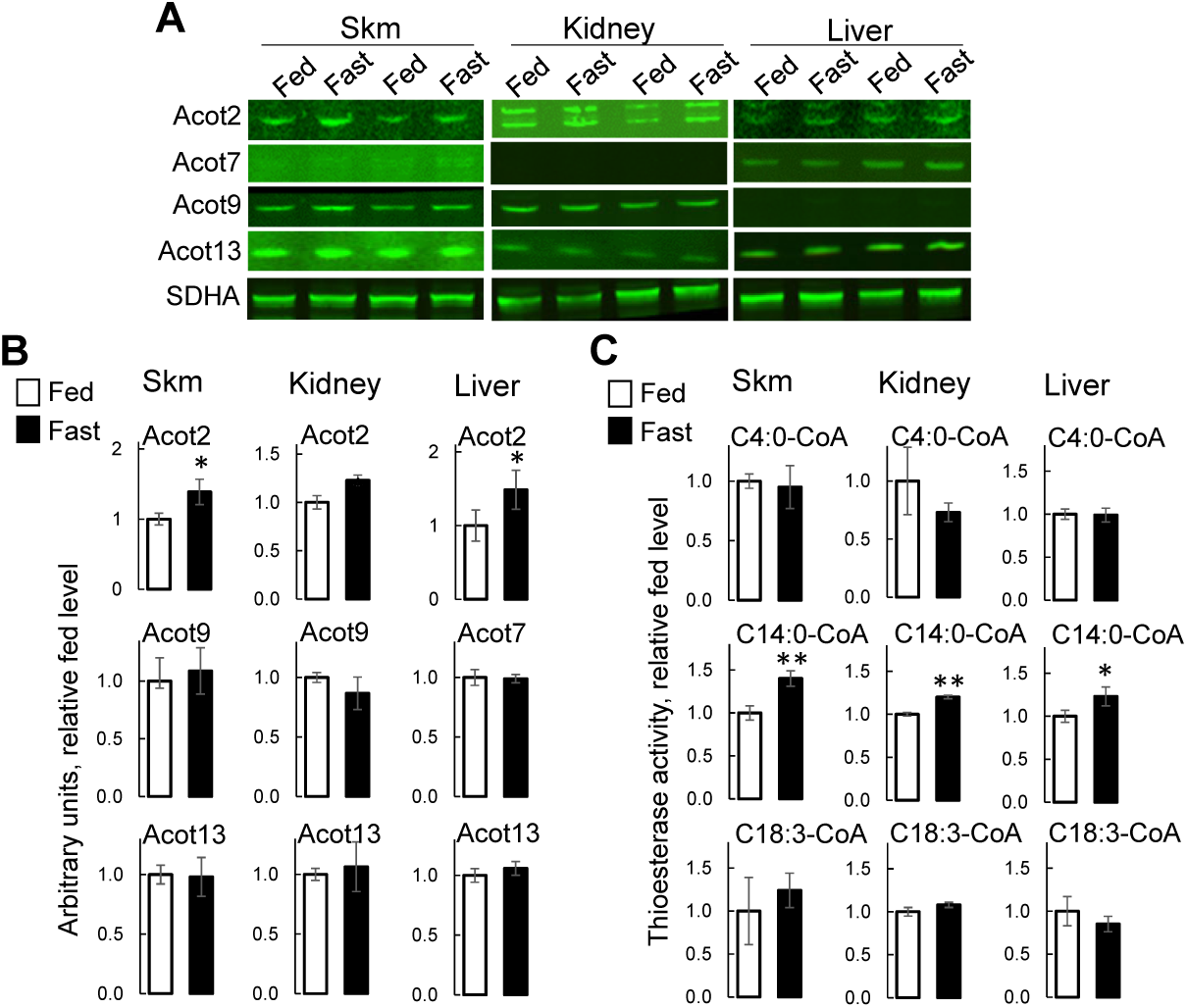
Fasting induces only small changes in mitochondrial Acot protein and thioesterase activity. Protein levels (**A, B**) and thioesterase activity (**C**) of Acots in mitochondria isolated from overnight fasted mice (∼9am tissue harvest) and *ad libitum* fed mice (∼5pm tissue harvest). Harvest times matched points of highest and lowest Acot2 mRNA expression (25). In **A** and **B**: Values: mean ± sem; n=4-5/condition. *: p=0.05, **p=0.02: unpaired t-test.

Beyond transcriptional regulation, there is some information on allosteric and other regulation of the Acots. However, this information is incomplete, and does not include potential allosteric regulators of Acot2, and also of Acot13 that would be relevant for its matrix localization. The activity of Acot1, which is ∼98% homologous to Acot2 at the amino acid level (29), was reported to not be regulated by CoA, though data were not shown (30). Using liver mitochondria overexpressing Acot2 (19) and hydrolyzing C14:0-CoA, we determined that Acot2 activity was insensitive to CoA up to 200 µM (**Fig5A**). Because Acot9 was found to be strongly inhibited by NADH (IC50 of ∼300 µM for C3:0-CoA and ∼500 µM for C14:0-CoA)(4), we determined the sensitivity of Acot2 and 13 activity to NADH and also to NAD^+^. Activity of neither Acot2 nor Acot13 was substantially changed by 1 mM NADH or 1 mM NAD^+^ (**Fig5A, B**). We also tested for sensitivity towards ADP and ATP and found that Acot2 and Acot13 activity was inhibited by ∼20% by 10 mM ADP or ATP (**Fig5A, B**).

**Figure 5.**
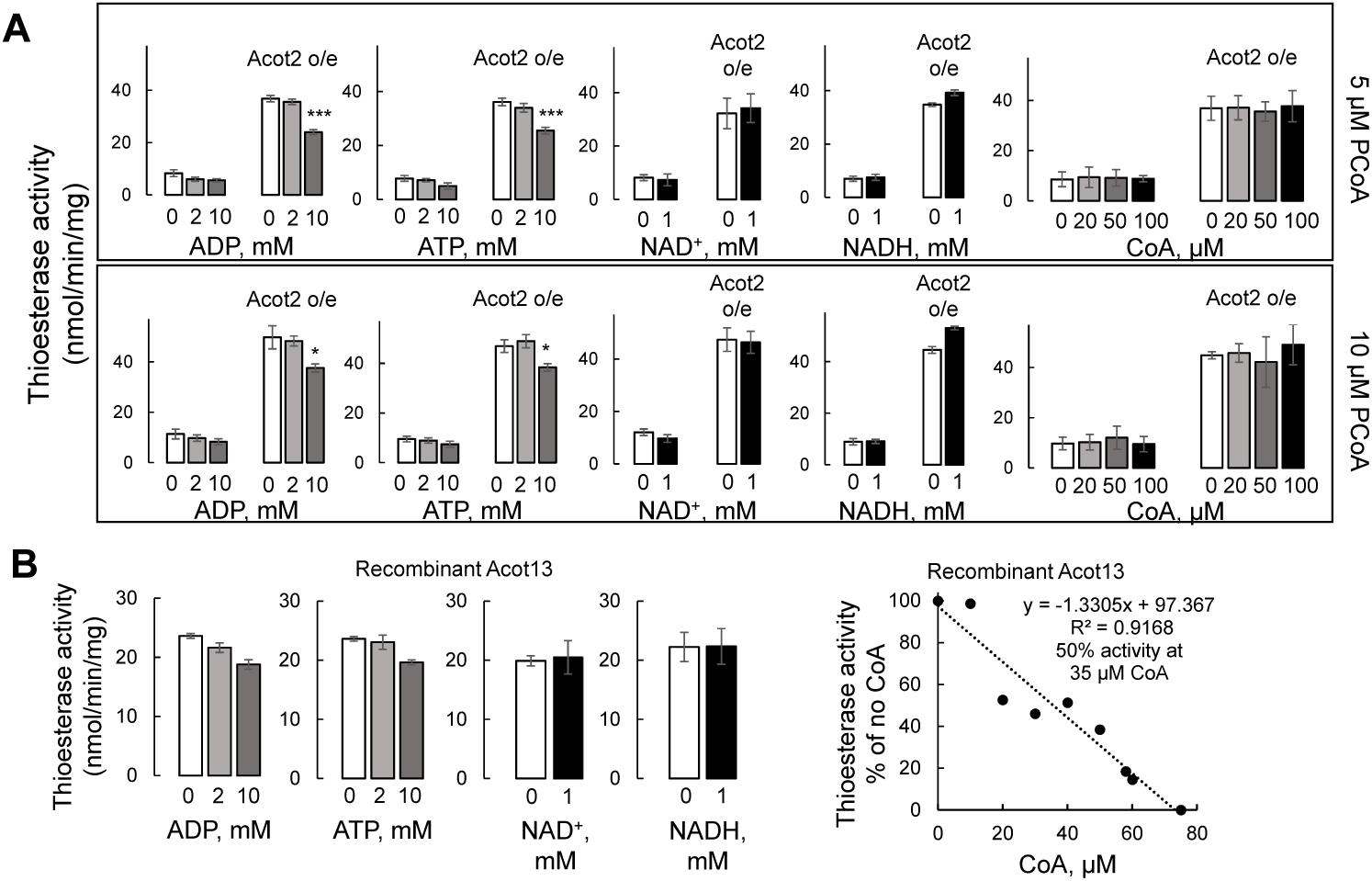
Distinct allosteric regulation of mitochondrial Acots. **Panel A**: Allosteric regulation of Acot2. Acot2-specific thioesterase activity towards 5 and 10 µM C16:0-CoA (palmitoyl-CoA: PCoA) in liver mitochondria without and with Acot2 overexpression. Values: mean±sem; n=4-6. *: p=0.02; ***: p<0.001; both p values are 10 mM *vs*. 0 mM ADP or ATP and 10 mM *vs*. 2 mM ADP or ATP; two-way ANOVA, Tukey post hoc tests. **B**: Allosteric regulation of recombinant Acot13, using C14:0-CoA (10 µM) as the substrate. Values: mean±sem; n=4 technical replicates. **C**: Immunoblot analysis of StarD2 expression in fractions from the mitochondrial isolation. Cytochrome c (cyto c) was used to mark mitochondria.

CoA has been reported to be a major regulator of Acot7 and Acot9 activity which were both inhibited by CoA (4,31). For Acot9, the [CoA] at which activity was inhibited by 50% (IC50) was ∼100 µM with C3:0-CoA as the substrate and ∼150 µM with C14:0-CoA as the substrate. Though it is difficult to accurately ascertain [CoA] within the mitochondrial matrix, especially since [CoA] is likely to be heterogeneous, inhibition of Acot activity by 100-150 µM CoA is likely to be physiologically relevant. To determine if Acot13, also a Type II Acot, is inhibited by CoA, we used recombinant mouse Acot13 (1). Acot13 activity towards C14:0-CoA was strongly inhibited by CoA with an IC50 of ∼35 µM (**Fig5B**).

Phosphotidylcholine transfer protein (PC-TP, also known as StARD2) interacts with Acot13 (32). Recombinant PC-TP almost doubled the Vmax of recombinant Acot13 (1,32); thus PC-TP could be considered as a regulator of Acot13. Because we found a significant fraction of Acot13 in the mitochondrial matrix, we asked if PC-TP was also present there. Using the protease protection approach we determined that PC-TP was associated with mitochondria (**Fig6A**), as reported (1), and also with the post-nuclear supernatant fraction of the 8000g spin, consistent with its presence in the cytosol (1). Mitochondria-associated PC-TP appeared to be fully cleaved by trypsin alone, analogous to OMM-localized Tom20 (**Fig6A**). Trypsin alone also cleaved Tim23 (IMM), and we had trouble obtaining conditions that would distinguish Tom20 and Tim23. We conclude that PC-TP localizes to the OMM, IMM and/or the intermembrane space. If it is in the matrix, it is in very low abundance; thus the majority of matrix Acot13 would not be influenced by PC-TP.

**Figure 6.**
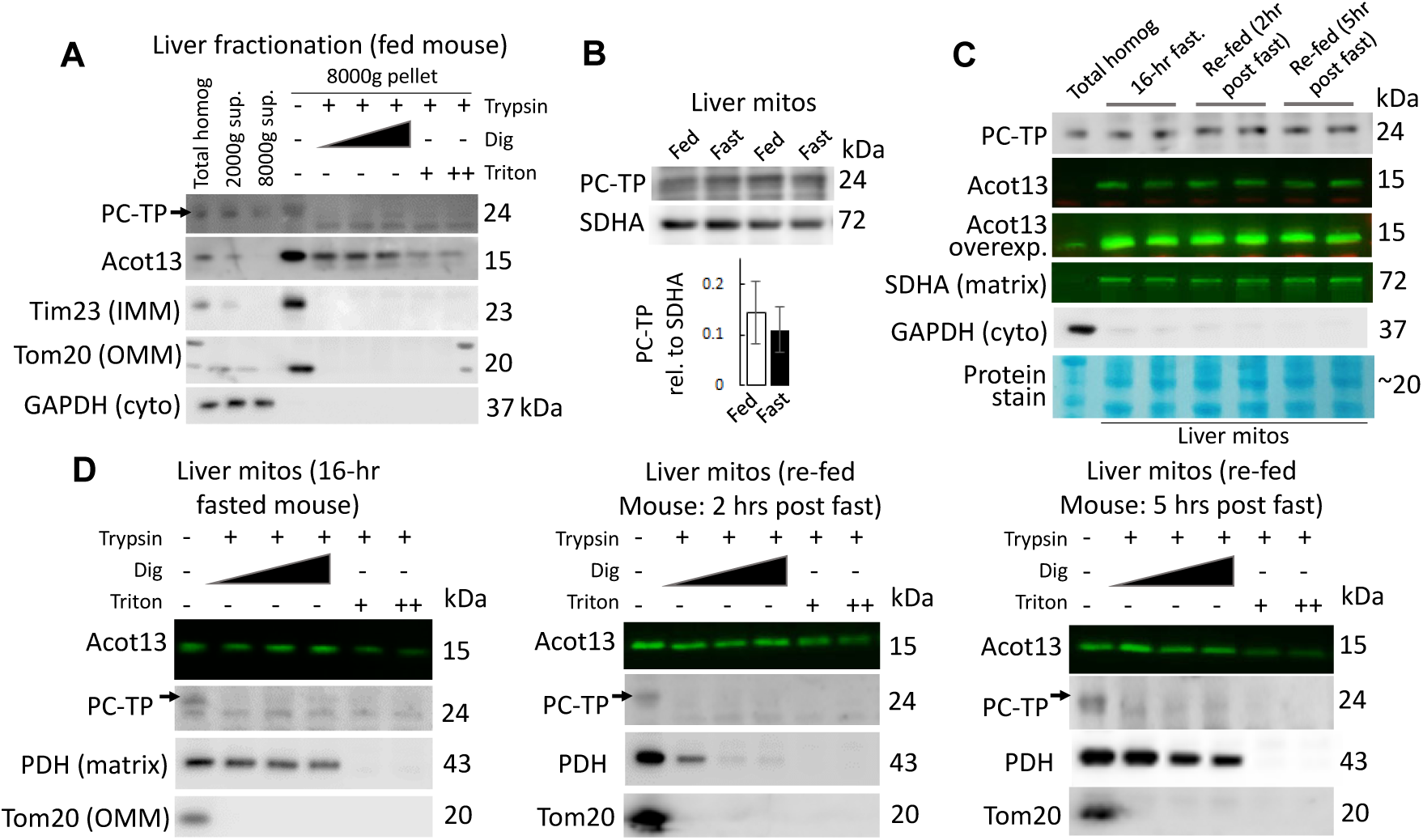
Regulation of Acot13 by PC-TP is only possible for the extra-matrix Acot13 pool and is unaffected by major nutrient perturbation. **A.** Liver fractionation and protease protection expertiment on the 8000g pellet fraction. GADPH was used as a cytoplasmic (cyto) marker. Sup.: supernatant. **B**. Mitochondrial fraction from liver obtained from *ad libitum* fed or overnight fasted mice. Bar chart: mean ± SEM, n=5 per condition. **C.** Mitochondrial (8000g pellet) fraction from liver obtained from mice fasted overnight, or fasted overnight then refed for 2 or 5 hours, and, **D.**, protease protection experiments from these mitochondria. In **C**, homogenate fraction is from a fasted condition and is used as a positive control for GADPH that is used as a cytoplasmic marker; mitochondrial fractions are not contaminated by cytoplasm. **Panels A, D**: Further details about protease protection experiments are available in the text describing Fig1.

The interaction of PC-TP with Acot13 in the liver was shown to increase during the early phase of re-feeding after mice were fasted overnight (7). Morover, PC-TP localization to mitochondria increased with PPAR agonist treatment (33). These findings prompted us to ask if the fraction of Acot13 available to be regulated by PC-TP depends on nutrient availability. We questioned, are fasting and refeeding associated with a redistribution of Acot13 away from mitochondria, was Acot13 associated with the mitochondrial membranes during refeeding, and did the association of PC-TP with mitochondria change? To answer these questions we fractioned the liver from fed mice, mice fasted overnight, or fasted then refed for 2 or 5 hours, then performed protease protection experiments on the 8000g pellet fraction. First, fasting did not influence the abundance of PC-TP associated with mitochondria (**Fig6B**). Comparing fasting to refeeding, we found no difference in the abundance of PC-TP, or Acot13, associated with mitochondria (**Fig6C**), and nor with the localization of mitochondrial Acot13 to the matrix and PC-TP to the OMM/IMS/IMM (**Fig6D**).

## Discussion

Here we undertook a comparison of mitochondrial thioesterases, in terms of activity, expression level and regulation, among mitochondria from different tissues in order to provide insight into why there are so many matrix-localized Acots. It was already suggested that several matrix Acots existed, some with an overlapping tissue expression and substrate specificity, raising the question of whether some matrix Acots are redundant. Alternatively, different matrix Acots could occupy distinct functional niches, defined by tissue specificity and/or regulation. These are important questions because matrix Acots are poised to regulate mitochondrial β-oxidation which is susceptible to overload with consequences on β-oxidation itself (5,34), on other matrix pathways and ultimately on other aspects of cell function. Thus understanding the characteristics of Acots, both individually and as a group, should provide insight into how Acots influence β-oxidation, and also how best to study the matrix Acots. However, distinguishing between the alternative hypotheses for why there are many matrix Acots has been precluded by the disbursed and incomplete information on Acots in the literature. The current study sought to fill key gaps and to provide easy comparisons among thioesterase activities and expression of Acots in mitochondria from several mouse tissues.

The main findings are summarized in **Figure 7** and are as follows. First, our study adds a new member to the matrix Acots, Acot13. This finding is important in its own right because of the substantial amount of existing data on the biological role of Acot13 (for review: (2,7)). Second, all mitochondria surveyed, except from liver, robustly expressed Acot2, Acot9 and Acot13. Liver, in contrast, expressed some Acot7 and Acot13 and very little Acot2. This pattern of mitochondrial Acot expression and activity across tissues highlights the potential importance of Acots in tissues such as kidney, skeletal muscle and heart, where Acots have been understudied. Third, Acot9 protein expression was previously shown to be very low in skeletal muscle and with a band shifted higher in molecular weight, and to be undetectable in heart (4). Here we show substantial thioesterase activity for C4:0-CoA in heart and skeletal muscle mitochondria. Acot9 is the only matrix Acot that hydrolyzes C4:0-CoA, and heart and skeletal muscle mitochondria each expressed Acot9 at a molecular weight similar to that in BAT and kidney where Acot9 was shown to be highly expressed (4)(present study). Thus we establish here that Acot9 is robustly expressed in heart and skeletal muscle, especially glycolytic skeletal muscle. Fourth, the most important source of regulation of the Acots, specifically the Type II Acots, was from CoA. In contrast, Acot2, a Type I Acot, was not regulated by CoA, and nor significantly by any of the potential allosteric regulators we tested, raising the possibility that Acot2 is constitutively active; we have provided an estimate for Acot2 activity in different tissues using a new mouse model of Acot2 loss. Altogether these observations reveal that, despite potential redundancy due to some overlap in tissue expression and substrate specificity (i.e., for Acot2 and Acot13), different matrix Acots largely occupy different functional niches determined mainly by regulation (Acot7, 9, 13 *vs*. Acot2) and substrate specificity (Acot9 *vs*. Acot2 and 13).

**Figure 7.**
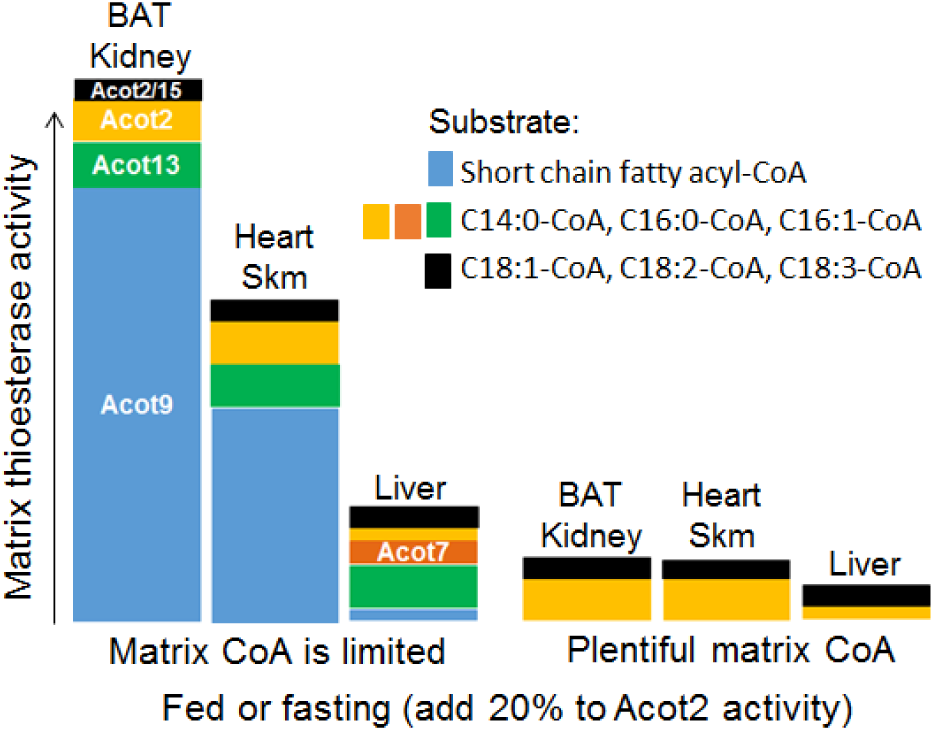
Summary of findings. Mitochondrial matrix Acots reside in different functional niches based primarily on substrate specificity and sensitivity to CoA inhibition. The height of the columns represents approximate thioesterase activity. The total height is the approximate total thioesterase activity for all substrates tested. Different colors represent thioesterase activity for a particular substrate or group of substrates, as indicated by the legend. The “Acot” member written inside the columns represents the Acot family member that would be responsible for the thioesterase activity represented by the color of the column. For each tissue, columns are shown for CoA-limiting (left grouping of columns) and CoA-replete conditions (right grouping of columns). Brown adipose tissue (BAT) and kidney mitochondria show the highest total potential thioesterase activity, contributed primarily by activity towards short-chain fatty acyl-CoAs, whereas liver mitochondria would have very low thioesterase activity, even under CoA-limited conditions. A CoA-limited condition would increase thioesterase activity for Acot7, 9 and 13 substrates, but not for Acot2 substrates. CoA was by far the strongest regulator of Acot9 and 13, and Acot2 was minimally regulated by multiple potential allosteric regulators (see Figure 5). Thus we propose a model in which Acot2 provides a constitutive overflow pathway for acyl-CoAs that are likely primarily destined for β-oxidation, whereas Acot7, 9 and 13 would provide an overflow pathway that depends on the local CoA concentration. Physiological activation of PPAR activity by fasting had only a small effect on Acot2 activity (and protein level), despite the substantial inducibility of Acot2 by PPAR agonists reported in the literature. See text for references. Skm: skeletal muscle.

Our study also highlights that, in contrast to the robust upregulation of Acot mRNA by pharmacologic induction of PPARs (1,15,17,25,27), fasting, a physiological stimulus of PPAR activity, had a minimal impact on mitochondrial Acot expression or activity. Thus, despite pharmacological approaches suggesting PPAR-induction of Acot mRNA as an important means of regulating Acots, regulation by CoA of Type II Acots may be quantitatively the most important physiological means of regulating the activity of matrix Acots (Type I or Type II).

Finally, because PC-TP has been shown to increase the activity of Acot13 (1), we determined if PC-TP could regulate matrix-localized Acot13. We found PC-TP to associate with mitochondria, as previously shown (1,33); most of the mitochondrial fraction was disrupted by trypsin alone and thus is likely on the OMM. This localization of PC-TP at mitochondria, and its abundance, were the same in liver from mice that were *ad libitum* fed, overnight fasted or refed after an overnight fast. Moreover, the abundance of Acot13 and its matrix-localization remained similar in all three conditions. These results are noteworthy because the amount of Acot13 that associates with PC-TP was higher in the liver of mice that were refed after an overnight fast (7); our findings suggest that this greater association of Acot13 and PC-TP involved the cytoplasmic pools of these proteins whereas the matrix pool of Acot13 was unperturbed. Collectively our data suggest a model in which there are 2 pools of Acot13 that are largely distinct in their association with PC-TP. A cytoplasmic Acot13 pool is available to interact with PC-TP, and the extent of this interaction is variable (7). A separate mitochondrial matrix pool of Acot13 is mostly or entirely unassociated with PC-TP.

### Localization of Acot13 to the mitochondrial matrix

Using two different proteases and mitochondria from three different tissues, we provide evidence that Acot13 localizes to the mitochondrial matrix, even though the prediction of an N-terminal MTS or an internal targeting sequence is low (∼30%). Most proteins that localize to mitochondria contain an N-terminal MTS that is cleaved, or, in fewer cases, an internal targeting sequence (10). However, some proteins that have been experimentally localized to mitochondria have low MTS predictions (e.g., MRPS22, MRPL42 and MRPL48) (12). Here we also show that, while an MTS prediction cutoff of 65% excludes most non-mitochondrial proteins, an MTS prediction below 65% has a high false negative rate. Thus, not only is there a precedent for mitochondrial matrix proteins devoid of an MTS, but a low MTS score is not strong evidence that a protein does not localize to mitochondria.

Two studies that documented the acetylated proteome of mouse liver mitochondria showed a significant acetylation of residue K13 of Acot13 that was dependent on the matrix deacetylase Sirt3 (35,36). Sirt3-dependent acetylation of Acot13 was also observed in heart mitochondria, though the specific residue was not indicated (37). Further analysis of these acetylated proteomes revealed that established OMM proteins with acetylated lysines did not have any that were Sirt3-dependent. These observations further support the matrix localization of Acot13.

### Regulation of matrix Acots

We found Acot13 to be strongly inhibited by CoA (Fig. 5), as reported for Acot7 and Acot9 (4,31). Though it is difficult to measure [CoA] in the matrix, the [CoA] that inhibits Acot9 and 13 (IC_50_ ∼35 µM for Acot13 and 100 µM for Acot9) is expected to fall within the physiological range, positioning these Acots as safety valves gated by CoA. Heart, skeletal muscle, kidney and BAT mitochondria substantially expressed Acot9 and 13 (Fig. 3) which, together, have a substrate specificity covering short- to long-chain acyl-CoAs. Thus the ability to expand the capacity of thioesterase activity in response to limiting CoA would apply across a wide range of fatty acyl-CoAs in the latter tissues. In contrast, liver mitochondria expressed minimal Acot9. Thus CoA gating would only occur in liver mitochondria for long-chain acyl-CoAs, via Acot13. Also, because of low Acot9 expression, the capacity to expand thioesterase activity in response to a CoA limitation would be much less in liver mitochondria.

Acot2 activity was only slightly inhibited by ADP and ATP, and was unaffected by the other potential allosteric regulators we tested including CoA, as was found for Acot1 (30). Thus, differently from Acot7, 9 and 13, Acot2 activity is not regulated by CoA. It is possible that acetylation regulates Acot2 activity because two acetylated lysine residues appeared in mitochondrial protein acetylation screens (35,36). Based on the crystal structure of human Acot2 (38), one of these lysines (K345 in human Acot2) is near the active site but pointed away from it. The other lysine (K104 in human Acot2) is in a loop that could be a flap over the active site; since acetylation imparts a negative charge, and CoA is negatively charged, K104 acetylation might alter substrate binding. Yet, the acetylation state of both lysine residues was insensitive to Sirt3 (35,36), decreasing the likelihood of regulation by acetylation as a means by which Acot2 activity is responsive to the matrix milieu. Acot2 may instead serve as a low capacity syphon for longer-chain fatty acyl-CoAs that runs independently of matrix metabolism.

### Considerations for evaluating the role of matrix Acots in β-oxidation

For all matrix Acots, the obvious competing pathway is β-oxidation because the fatty acyl-CoA dehydrogenases could compete directly for fatty acyl-CoA substrates. Competition would be on the basis of substrate specificity, and also Km, the values of which for Acot2 and 13 are similar to those of the long-chain acyl-CoA dehydrogenase reaction for those substrates (39). Because short-chain methyl-branched acyl-CoAs are also substrates for Acot9, a role in amino acid oxidation was also proposed (4) but has not yet been tested.

To determine the potential impact of matrix thioesterase activity on β-oxidation, values of thioesterase activity (Fig. 3) can be compared to β-oxidation rates. The heart is a useful test case because there are substantial data on palmitate oxidation using ^3^H(9,10)-palmitate which directly assesses flux through the acyl-CoA dehydrogenases. In working heart preparations, palmitate oxidation rate is ∼15 nmol/min/mg mito protein (based on ∼1200 nmol/min/dry heart weight and mitochondria at 2% of wet heart weight) (40-42). In heart mitochondria, maximal thioesterase activity towards C16:0-CoA was not much less, at ∼6 nmol/min/mg. If activities for C14:0-CoA, C4:0- and C8-CoA are also taken into account, and considering lower β-oxidation rates in tissues other than heart, thioesterase activity could theoretically outstrip palmitate oxidation. Is this actually possible?

Direct evidence that a β-oxidation substrate can be hydrolyzed to fatty acid, and thus that β-oxidation competes with thioesterase activity, was shown by supplying mitochondria with ^14^C-palmitoylcarnitine (20 µM) then measuring ^14^C-palmitate production by thin layer chromatography (16,20). Palmitate was produced at ∼0.03 nmol/min/mg mito protein in skeletal muscle mitochondria (20) and 0.5 nmol/min/mg mito protein in heart mitochondria (16). These values are 1-2 orders of magnitude lower than the maximal thioesterase activity for palmitoyl-CoA, suggesting that matrix thioesterases offer far less competition for β-oxidation then is predicted by maximal thioesterase activities and their Km values. Possible explanations for this discrepancy include unknown regulators of the Acots and an *in vivo* limitation on access to the Acots by substrates, due either to tight substrate channeling within β-oxidation (43), or because the Acots are located apart from the β-oxidation machinery. Also, the CoA dependence of Acot13 means that, depending on CoA abundance, CoA inhibition of Acot13 could remove a competitor of palmitate oxidation. Likewise, removal of short-chain fatty acyl-CoA from the β-oxidation of palmitate would also depend on the extent of CoA inhibition of Acot9. Thus several factors independently or together would decrease the competition between Acots and β-oxidation, and at least one factor, CoA availability, would alter the extent of the competition between these pathways. It should be noted that all of the tissues surveyed here expressed very little Acot activity towards C18:0-CoA or unsaturated C18 CoA esters. The latter are the major substrates for Acot15. Thus, Acot15’s ability to compete with β-oxidation may be minimal.

### Possible functional niches for matrix Acots in the context of β-oxidation

Evidence that Type II matrix Acots have a functional role that depends on CoA abundance is in line with the idea that matrix Acots mitigate a CoA limitation, as originally proposed by Himms-Hagen and Harper (44). However, the CoA independence of Acot2 activity implies an additional role for matrix Acots. This role might be as a minimally regulated syphon that removes long-chain fatty acyl-CoA from β-oxidation.

An ability of matrix Acots to compete with β-oxidation implies that matrix processes can exert control on β-oxidation, a concept that conflicts with the generally accepted model that the main control point for β-oxidation is the entry of activated long-chain fatty acids into mitochondria at CPT1 (carnitine palmitoyl-transferase-1) at the OMM. However, recent experimental and modeling studies show that control of β-oxidation can be partly or substantially taken over by the β-oxidation reactions themselves (5,34). This shift in control occurred at high concentrations of long-chain fatty acyl-CoA and lead to the accumulation of intermediates and partial depletion of CoA (34). This so-called β-oxidation overload was attributed to substrate competition at the β-oxidation enzymes because they have activity towards multiple substrates (34). The possibility that β-oxidation enzymes exert control over β-oxidation was also suggested by measurements of β-oxidation intermediates (for review: (45)). More recent modeling studies indicate that, at high C16:0-CoA concentration, control of β-oxidation shifts to the C4-C6 reactions of MCKAT (medium-chain ketoacyl-CoA thiolase) (5). This potential for overload, which could theoretically halt β-oxidation (5), suggests the utility of the Acots to mitigate gridlock at MCKAT and to prevent CoA depletion.

Models of β-oxidation do not include Acot reactions (5,34). The present study facilitates the incorporation of Acots into these, which could help to shed further light on the extent of the influence of Acots on β-oxidation, including in conditions of potential substrate overload.

**In conclusion**, the present study has determined that Acot7, Acot9 and Acot13 reside in the mitochondrial matrix, joining Acot2 and Acot15. Furthermore, we report that these matrix Acots are not functionally redundant but, rather, are specialized according to substrate specificity and sensitivity to CoA inhibition. These distinct yet potentially complementary roles of the matrix Acots predict that their functional relevance will increase with decreasing CoA availability to ensure that CoA does not become limiting for β-oxidation and also for branched chain amino acid catabolism. However, our study also reveals that Acot2 is minimally influenced by factors within the matrix including CoA. Thus Acot2 may be a constitutively active matrix thioesterase and have a role beyond mitigating CoA limitation.

## MATERIALS AND METHODS

Details are provided in **Supporting Information.**

### Reagents

Acyl-CoA esters were purchased from Avanti Polar Lipids, and used for no longer than 2 months after dilution. Most other reagents were purchased from Sigma-Aldrich with the exception of antibodies and recombinant Acot13.

### Control and Acot2 knockout Mice

Ten-14 wk old male were used in accordance with protocol #01307 approved by the Thomas Jefferson University Institutional Animal Care and Use Committee. C57BL/6J mice or whole-body Acot2 knockout mice (Acot2^-/-^) were generated from our breeding colony. The strategy used to generate mice with floxed Acot2 alleles (Acot2^fl/fl^) and Acot2^-/-^ mice is described in **Fig3B** and **Supporting Information**. Mice were maintained on a 12-12h light-dark cycle (lights on: 7am – 7pm), and *ad libitum* fed a standard diet (LabDiet 5001, Purina). Some mice were fasted overnight (7pm – 9am). Mitochondria from whole-body Acot13 knockout mice (7) were isolated in the lab of Dr. David Cohen.

### Expression of recombinant Acot2 in mouse liver

Adenovirus harboring mouse Acot2 cDNA or empty vector control (2X10^9^ pfu/mouse; as in (19)) was diluted in saline then administered to mice via the tail vein, as in (19). Liver mitochondria were isolated seven days post injection. Localization of Acot2 to mitochondria and not to cytosol was previously confirmed, as well as the appropriate substrate specificity for Acot2 (19).

### Isolation of mitochondria from mouse tissues

Mice were sacrificed by cervical dislocation at ∼9am. All media were ice cold, and the procedure was done on ice or at 4 °C. Protein of the final resuspended pellets was determined by bicinchoninic assay (BCA; Life Technologies, Carlsbad, CA, USA). Detailed protocols are provided in **Supporting Information**.

### Thioesterase activity

Thioesterase activity was measured in isolated mitochondria (19) or recombinant Acot13 protein (1), essentially in as (19).

### qPCR and immunoblotting

Information about primers and antibodies is provided in **Supporting Information**.

### Protease protection experiments

Reactions were run for 15 minutes, at room temperature, in a total volume of 40 ul of MAS, using 2.5 mg/ml of mitochondria. During the assay we tested different concentrations of trypsin: 0.068 mg/ml, 0.102 mg/ml, 0.152 mg/ml and 0.204 mg/ml of final concentration for each reaction; as well as proteinase K: 0.01 mg/ml or 0.02 mg/ml of final concentration. Digitonin was added at final concentration 1 μg/μl, 1.25 μg/μl, or 1.5 μg/μl, and Triton at a final concentration of 1% or 2%. Reactions were stopped by 10 min incubation at room temperature with 3.25 mM PMSF.

### Statistical analysis

Bar charts show the mean and standard error of the mean. Statistical analysis was done using either a Student’s t-test or analysis of variance followed by Tukey post hoc tests, as appropriate; details are provided in the figures and figure legends. SigmaPlot was used to perform the statistical analysis. Further details of the statistical are described in the text and figure legends.

## Acknowledgements

This work was funded by the American Heart Association 13SDG14380008 and NIH R01 DK109100 (to E.L.S.), and Thomas Jefferson University start-up funded (to E.L.S.). We thank Megan Roche and Cynthia Moffat for contributions to preliminary experiments.

