## Supporting Information for "Multiple acyl-CoA thioesterases occupy distinct functional niches within the mitochondrial matrix"

**Supporting Information Methods:**

**Generation of mice with floxed Acot2 alleles and with whole-body deletion of Acot2:**

The general targeting strategy was to flank exon 2 with loxP sites. To this end, a 7.96 Kb region used to construct the targeting vector was first subcloned from a positively identified C57BL/6 BAC clone (RP23: 326I12). The region was designed such that the short homology arm (SA) extended ~2.01 Kb 3' to the Neo cassette and the long homology arm (LA) ended 5' to exon 2, and was 5.40 Kb long. The loxP/FRT flanked Neo cassette was inserted 195 bp downstream of exon 2. The single LoxP site, containing engineered Afl II, Avr II and Nde I sites for southern blot analysis, was inserted 159 bp upstream of exon 2. Thus the target region was 556 bp and included exon 2. The targeting vector was confirmed by restriction analysis after each modification step, and was linearized using Not I for electroporation into C57Bl/6N embryonic stem (ES) cells. An initial screen identified potential positive clones. These were further screened for SA and LA integration and also for the thrid LoxP site. Screening was by PCR and also by Southern blotting as a secondary analysis. Several clones were confirmed as correctly targeted and recommended for injection into Balb/c blastocysts. Resulting chimeras with a high percentage of black coat color were mated to wild-type C57BL/6N mice to generate F1 heterozygous offspring. Tail DNA was analyzed from pups with black coat color; a PCR was used to detect the presence of the distal LoxP site and also the SA. The Frt-flanked Neo cassette was removed by crossing heterozygous floxed Acot2 mice with FLPo Deleter mice. Heterozygotes Neo-deleted mice ($Acot2^{fl/wt}$) were then bred with ubiquitously expressed Cre (Ella-Cre) mice to generate germ-line whole-body Acot2 knockout mice ($Acot2^{-/-}$).

**Isolation of mitochondria from mouse tissues:**

Skeletal muscle mitochondria: Isolation of skeletal muscle mitochondria was performed essentially as described in (2). All steps were performed on ice or at 4°C. Skeletal muscle from all limbs was dissected and placed in skeletal muscle medium (SKM: 140 mm KCl, 20 mm, HEPES, 5 mm $MgCl_2$, 1 mm EGTA, pH 7.0). Muscle was cleaned of connective tissue and fat, minced, and placed in 15 volumes of homogenizing medium (SKHM: SKM with 1 mm ATP and 1% BSA (w/v)) containing two units of protease (from Bacillus licheniformis, Sigma P5380) per g of muscle wet weight. Tissue was homogenized using a glass/Teflon Potter-Elvehjem homogenizer (500 rpm, 12 passes), then fractionated by centrifugation at 600*g* (10 min), and the supernatant was collected and spun at 10,000*g* (10 min). The pellet was resuspended in SKM then incubated on ice for 5 min (myofibrillar repolymerization). Samples were spun at 600*g* (10 min), then supernatant was filtered then spun at 10000*g* (10 min). The final pellet was resuspended in SKM to achieve a concentration of ~ 25 mg/ml.

Liver mitochondria: Liver mitochondria were isolated as described (3). All steps were performed on ice or at 4°C. Liver was dissected, washed in liver medium (LM; 250 mM sucrose, 10 mM Tris-HCl, 0.1 mM EGTA, pH 7.4), minced, then suspended in LM+0.5% defatted BSA in a glass/Teflon Potter-Elvehjem homogenizer and homogenized (500 rpm, 12 passes). Samples were centrifuged (10 min, 600*g*), then again (10 min, 600*g*), then for 10 min at 7000*g*. The pellet was resuspended in LM then washed twice. The final pellet was resuspended in LM to have a protein concentration of ~25 mg/ml.

Heart and kidney mitochondria: Heart mitochondria were isolated as described in (4), and kidney mitochondria followed the same procedure. All steps were performed on ice or at 4°C. Kidney or heart was dissected, washed in heart/kidney isolation buffer (HKB: 225 mM mannitol, 75 mM sucrose, 20 mM HEPES, 0.1 mM EGTA, pH 7.4), minced, then suspended in HKB+0.5% defatted BSA in a glass/Teflon Potter-Elvehjem homogenizer and homogenized (350 rpm, 10 passes). Samples were centrifuged (5 min, 500*g*), then (10 min, 9000*g*), washed the pellet and centrifuge again (10 min, 9000*g*). The final pellet was resuspended in HKB in a volume that resulted in a protein concentration of ~25 mg/ml.

Brown adipose tissue (BAT) mitochondria: BAT mitochondria were isolated as in (5). All steps were performed on ice or at 4°C. BAT was dissected, washed in BAT isolation buffer (BATB: 250 mM sucrose, pH 7.4), minced, then suspended in BATB+0.2% defatted BSA in a glass/Teflon Potter-Elvehjem homogenizer and homogenized (350 rpm, 10 passes). Samples were centrifuged (10 min, 8500*g*), then (10 min, 500*g*), and again (10 min, 8500*g*). The final pellet was resuspended in BATB in a volume for a protein concentration of ~25 mg/ml.

**Immunoblotting**

During the experiments we used custom antibodies and commercially available once summarized in Table below. Samples were loaded onto an 10% or 12% polyacrylamide gel (for Licor-conjugated secondary antibodies) or a 4-12% gel (chemiluminescence (ECL)-conjugated secondaries), electrophoresed, transferred onto nitrocellulose and blocked with Odyssey blocking buffer (LI-COR Biosciences, Lincoln, NE, USA) diluted in PBS or with TBS-T plus 3% BSA when ECL was used for detection. Primary antibodies were diluted in TBS-T. Secondary antibodies (LI-

COR anti-MS 92632212 and anti-Rb 92632213) were diluted 1:20000 in 5% skin milk or in TBS-T for ECL (Thermo Fisher anti-MS 31473 and anti-Rb 31460). Protein bands were visualized using the LI-COR Odyssey3000 system or after application of ECL (SuperSignal™ West Dura Extended Duration, Invitrogen) followed by visualization (Kodak Image Station; 440CF; CareStream MI software). Quantification and normalization are explained in Results.

**Antibodies**

| Antibody | Source | Dilution |
|---|---|---|
| Acot2 | Stefan Alexson, Karolinska Institute (e.g.[Moffat, 2014]) | 1:1000 |
| Acot9 | Stefan Alexson, Karolinska Institute [Tillander, 2009] | 1:1000 |
| Acot7 | Michael Wolfgang, Johns Hopkins University [Ellis, 2015] | 1:1000 |
| Acot13 | David Cohen, Weill Cornell (e.g [Wei, 2009]) | 1:1000 |
| StarD2 (PC-TP) | David Cohen, Weill Cornell (e.g. [Wei, 2009]) | 1:1000 |
| Complex I (39 kDa subunit) | Abcam Cat#Ab14613 | 1:1000 |
| Complex II (SDHA subunit) | Abcam Cat#Ab14715 | 1:1000 |
| Complex III (Core 1 subunit) | Abcam Cat#Ab14745 | 1:1000 |
| GAPDH | Sigma; Cat# MAB374 | 1:10000 |
| Cytochrome c | BD Pharmingen; Cat#44411 | 1:1000 |
| Grb2 | Santa Cruz; Cat#C-73 | 1:1000 |
| PDH | Abcam; Cat#110330 | 1:1000 |
| TOM20 | Santa Cruz; cat#sc-11415 (no longer available) | 1:1000 |
| TIM23 | DB Bioscience; Cat#51344 | 1:1000 |

**qPCR**

Total RNA was extracted from tissue using Trizol® (Invitrogen, Carlsbad, CA), then treated with RQ1 DNase (Promega, Madison, WI, USA) at 37$^{o}$C for 30 mins. RNA concentration was measuring using a Qubit® Fluorometer (Invitrogen, Carlsbad, CA). RNA was reverse transcribed using oligo(dT)$_{20}$ primers and SuperScript III (Invitrogen, Carlsbad, CA). Primers were designed using Eurofins Primer Design Tool, and checked for specificity and efficiency. Custom oligos were purchased from Eurofins MGW Operon (Huntsville, AL). Primers sequences are shown below. qPCR reactions were performed using ITaq SYBR green Supermix with ROX (BIO-RAD, Hercules, CA) in 20 µl reactions (20 ng cDNA/reactions), and using an Eppendorf Mastercycler® ep realplex. The $\Delta\Delta$Ct was used to calculate transcript levels relative to β-actin.

**qPCR Primers**

| Primers | Sequence |
|---|---|
| Acot2 | Forward: TGGGAACACCATCTCCTACAA<br>Reverse: CCACGACATCCAAGAGACCA |
| Acot7 | Forward: ATCAGCACGCGGCACTGTAA<br>Reverse: TTGGTACCTGTGAGGATGTTCTCC |
| Acot9 | Forward: AGGTGGGAACCAAGATTCAG<br>Reverse: TTCAAGGTCCAAAGCCGTATC |
| Acot13 | Forward: AGCCCAGACTCTTGCTTTG<br>Reverse: TAGTTTCTCAGGAGCAGCC |
| Acot15 | Forward: TCAGACAAACAGGACTGGC<br>Reverse: ACCCTCCGTGAGCAAAC |
| Beta-actin | Forward: CAACACCCCAGCCATG<br>Reverse: GTCACGCACGATTTCCC |

**Skeletal muscle homogenates**

Skeletal muscles were rapidly dissected then frozen in liquid nitrogen. Muscles were suspended in RIPA lysis buffer (150 mM NaCl, 1% NP-40, 0.5% Na-deoxycholate, 0.1% SDS, pH 8.0), minced with scissors, then homogenized with the tube maintained on ice. Samples were cleared at 18000*g*, 4ºC. Protein concentration was determined by BCA.
